# Calorie Restriction modulates beta cell IP_3_R activity to regulate Ca^2+^ homeostasis and cell network connectivity

**DOI:** 10.1101/2025.09.01.673520

**Authors:** Johannes Pfabe, Cristiane dos Santos, Melanie Cutler, Christopher Acree, Aliyah Habashy, Amanda Cambraia, Guy Perkins, Mark H Ellisman, Marjan Slak Rupnik, Rafael Arrojo e Drigo

**Author notes:** Universidade de Campinas, Departamento de Bioquímica e Biologia Tecidual, Campinas, São Paulo, Brazil (Cristiane dos Santos).

## Abstract

Calorie restriction (CR) promotes beta cell longevity by regulating cell identity, organelle and protein homeostasis, and metabolism pathways. CR beta cells have higher cAMP levels and mitochondria with an elevated potential to generate ATP. However, CR beta cells have reduced insulin secretion due to increased peripheral insulin sensitivity. How CR impacts beta cell Ca^2+^ homeostasis to regulate beta cell insulin release remains unknown. We investigated this question using acute pancreatic tissue slices prepared from ad-libitum (AL) or CR mice loaded with a low affinity Ca^2+^ indicator and recorded cytosolic Ca^2+^ gradients with fast confocal imaging. We exposed these slices to increasing glucose concentrations and applied our semi-automatic analysis pipeline to detect thousands of individual beta cells followed by identification of individual Ca^2+^ spiking events. We observed that CR beta cells have fast short-amplitude Ca^2+^ oscillations that correlate with largely disconnected beta cell networks across the islet. Using acetylcholine stimulation, we found that faster IP_3_R-driven Ca^2+^ oscillations linked to higher cytosolic cAMP levels protect beta cells against acute depletion of ER Ca^2+^ stress. Therefore, this study provides new mechanistic insight into adaptation of beta cell and of beta cell networks to CR interventions.

**Article highlights:**

- Beta cells from calorie restricted (CR) mice have decreased insulin release, however the mechanisms underlying this adaptive response remain unknown.
- CR beta cells have elevated basal cytosolic cAMP ([cAMP]_cyt_) compared to beta cells in control ad libitum fed (AL) mice, and they operate with faster and shorter cytosolic Ca^2+^ oscillations.
- While AL beta cells form interconnected activity networks, CR beta cells are largely disconnected and fire more independently of each other.
- Islets of CR mice can sustain prolonged activity during ER stressing conditions due to elevated IP_3_R activity and improved Ca^2+^ homeostasis.

**Why did we undertake this study?:** We have previously shown that calorie restriction (CR) promotes beta cell longevity by enhancing beta cell identity and organelle homeostasis mechanisms. This long-lived phenotype correlated with the onset of enhanced peripheral insulin sensitivity and reduced beta cell insulin release in vivo despite higher cAMP levels and increased potential for mitochondrial ATP generation. However, the mechanisms underlying the reduced cell insulin release phenotype of CR beta cells remains unknown. Therefore, we investigated the underlying Ca^2+^ homeostasis mechanisms regulating insulin release in AL and CR beta cells.

**What is the specific question(s) we wanted to answer?:** We were interested in determining what are the cell Ca^2+^ activity patterns during basal and glucose-stimulated conditions in AL and CR beta cells. In addition, we also investigated how CR beta cells respond to epinephrine inhibition and supra-stimulatory concentrations of acetylcholine (ACh), which drive acute beta cell stress by disrupting normal cAMP and ER Ca^2+^ signaling, respectively. Finally, we investigate whether CR beta cells formed more interconnected beta cell networks driven by changes in Ca^2+^ activity patterns.

**What did we find?:** We found that CR beta cells are more active with significantly higher rates of Ca^2+^ oscillation at basal and high glucose concentrations. In fact, CR beta cells have shorter inter-Ca^2+^ event intervals that are more resistant to depletion of cAMP by epinephrine application. In contrast, stimulation of IP_3_R activity (to force depletion of ER Ca^2+^ stores) by supraphysiological ACh concentrations revealed that CR beta cells were able to sustain a prolonged Ca^2+^ activity versus AL beta cells. Surprisingly, this enhanced beta cell activity profile reduced beta cell activity network connectivity.

**What are the implications of our findings?:** Our work demonstrates that CR beta cells have higher baseline and glucose-stimulated Ca^2+^ activity due to higher cAMP levels. These cells also have dominant IP_3_R activity that grants improved ER Ca^2+^ homeostasis and significantly reduces beta cell network connectivity to tone down insulin secretion. These studies provide a mechanistic understanding of how beta cells adapt to CR and to CR-associated enhanced insulin sensitivity.

## Introduction

Ca^2+^ is being recognized as an important second messenger for insulin exocytosis since the late 1970s (1). The discovery of the excitability of beta cells (2) led to the classical model of Ca^2+^-mediated insulin release. In the current consensus model of insulin release (3), glucose gets taken up by the cell through glucose transporters and is metabolized in the mitochondrium yielding ATP. The rise of [ATP] triggers the closure of K_ATP_-channels leading to depolarization of the cell followed by a Ca^2+^ influx from the extracellular space through L-type Ca^2+^-channels. Insulin vesicles are then released via Ca^2+^-triggered exocytosis. Recent research challenges this model (4) advocating that the glucose metabolism plays a more important role than previously thought. Moreover, the influence of Ca^2+^ stored in the endoplasmic reticulum (ER), especially with focus on ryanodine receptors (RyR) and IP_3_-receptors (IP_3_R) (5), add more complexity to the model.

Calorie restriction (CR) is a dietary approach that improves several cardiometabolic indexes and can lead to increased organismal longevity and improved health span, from nematodes to mice to monkeys and to humans (6-11). From a mechanistic standpoint, CR works by reducing overall energy intake, which decreases metabolic stress by improving nutrient homeostasis and metabolism, and reducing the need for sustained insulin signaling pathways (12). In the pancreas, we and others have shown that CR modulates pancreatic beta cell gap junction connectivity and normalizes Ca^2+^ homeostasis in obese mice (13), enhances beta cell identity and survival in lean and obese mice (14), and promotes organelle and protein homeostasis while limiting insulin release levels in young adult mice (15). Together, these pathways work together with CR-mediated increase in peripheral insulin sensitivity to reduce the metabolic load on beta cells, thus decreasing beta cell turnover and enhancing beta cell longevity via protection from age-associated damage (15).

CR modulates the abundance of several secondary messenger molecules and pathways, including cyclic AMP (cAMP) and Ca^2+^ (16; 17). Using single cell sequencing and imaging mass spectrometry, we demonstrated that CR beta cells have increased basal islet cAMP levels and upregulation of several genes involved in ER structure-function (protein folding and degradation, and Ca^2+^ signaling) (15). In beta cells, both cAMP and Ca^2+^ signaling pathways converge to drive the amplification of glucose-induced insulin release via activation of protein kinase A (PKA) or Epac2 (18). Therefore, we hypothesized that the observed increase in basal cAMP levels in CR beta cells would lead to increased Ca^2+^ release pathways from the ER via ryanodine (RyR) or IP_3_-receptors (IP_3_Rs) and modulated beta cell insulin release dynamics.

We tested this hypothesis by exposing FVB mice to 20% CR for 8-weeks, which is sufficient to improve glucose homeostasis and reduce beta cell insulin release in vivo (15). We monitored beta cell Ca^2+^ homeostasis and beta cell activation dynamics using high speed confocal microscopy of in situ beta cells in acute pancreatic slices exposed to a glucose ramp and cAMP and ER Ca^2+^ modulators epinephrine (Epi) or acetylcholine (ACh), respectively. We found that CR beta cells have faster and shorter Ca^2+^ events, thus indicating that CR beta cells have finer control of insulin release versus control AL cells. Moreover, treatment of CR beta cells with epinephrine (to dampen cytosolic cAMP levels) or acetylcholine (to drive ER IP_3_R Ca^2+^ release) revealed that CR beta cells can sustain prolonged and faster Ca^2+^ oscillations, whereas AL beta cells rapidly exhaust their intracellular Ca^2+^ stores. None of these changes were explained by changes in ER ultrastructure or mitochondria-ER connectivity. These data indicate that CR beta cells can sustain normal function during stressful situations due to enhanced Ca^2+^ homeostasis mechanisms that are independent of changes to ER anatomy.

Finally, CR beta cells have mostly disconnected activity networks, and islet beta cell collectives have largely independent activity patterns. Together, our data demonstrate that CR activates Ca^2+^ spiking patterns in pancreatic beta cells that are primarily influenced by higher levels of cAMP. CR beta cells have faster Ca^2+^ oscillations due to a higher baseline of ER Ca^2+^ release and faster recovery to epinephrine-mediated cAMP depletion. Furthermore, CR beta cells have a higher capacity to endure high ER Ca^2+^ release demands. This work provides a mechanistic explanation regarding how beta cells adapt to CR to reduce insulin secretion.

## Research Design and Methods

### Animals and diet manipulation

All animal experimentation was approved by the Institutional Animal Care and Use Committee at Vanderbilt University (IACUC protocol M2000086). Mice were maintained in rooms with an average temperature of 23°C and with a 12h light and 12h dark cycle. For all our studies, we purchased 3-week-old FVB male mice from Jackson Labs (JAX, Connecticut, strain number 1800). CR feeding was performed as previously established by us (15). Briefly, mice were housed and acclimated using our standard study chow (Lab Diets, 5053) for 5 weeks prior to any experimentation. Next, we monitored the daily food consumption of each cage during the last week of the adaptation window to determine a food consumption baseline. At 8 weeks of age, mice were weighted and randomly assigned to AL or CR groups; mice in the CR group received 20% less daily grams of chow in relation to the AL group. Mice were kept on AL or CR diet for 8-weeks. Weight was monitored daily until the end of the study and mice were euthanized for pancreatic slice preparation at 16 weeks of age.

### Preparation of acute live pancreatic slices

We prepared acute pancreatic tissue slices as described previously (19). Briefly explained, we killed the mice by decapitation after isoflurane treatment. Following, we performed a laparotomy to access the pancreas, followed by injection of 1.9% low-melting point agarose dissolved in extracellular solution (ECS, [containing in mM: 125 NaCl, 10 NaHCO_3_, 10 HEPES, 6 lactic acid, 3 myo-inositol, 2.5 KCl, 2 Na-pyruvate, 2 CaCl_2_, 1.25 NaH_2_PO_4_, 1 MgCl_2_, 0.25 ascorbic acid; adjusted to pH 7.4 by adding NaOH; supplemented with 6 mM glucose if not stated elsewise]) into the common bile duct. Immediately after injection, we poured ice cold ECS over the exposed pancreas to let the agarose solidify. Next, we extracted the pancreas, embedded the tissue in agarose, and produced tissue slices of 140 µm thickness using a Leica VT1200S vibratome. In the end, we incubated the slices in ECS containing 6.865 µM Calbryte 520 AM, a low affinity chemical indicator for cytosolic Ca^2+^, and 0.0075 % Pluronic F-127 for 60 min at room temperature on an orbital shaker.

### Live Ca^2+^ imaging of mouse pancreatic slices

For the imaging, we put the slices into a low-volume chamber constantly perfused with ECS at 37°C. We used a confocal laser scanning microscope (Leica Stellaris X5) with a 20x/0.95 objective to image the slices. We excited the Ca^2+^ indicator with a white laser line set to 494 nm. Emitted light was acquired using a HyD detector tuned for collection between 500 nm and 700 nm in the photon counting mode. We recorded XYT-series with a frequency of 20 Hz and a resolution of 256 x 256 pixels. For the change of treatment conditions, we switched to solutions with different glucose concentrations, which we also combined with pharmacological compounds (Epi, ACh). We prepared slices from at least three mice per diet group and imaged at least three islets slices per mouse for each experiment described in this paper.

### Analysis of beta cell cytosolic Ca^2+^ patterns

We extracted the data from the raw imaging files as described previously (5). Briefly, we processed the raw data by segmenting a statistical image calculated from the mean pixel intensity plus the highest percentile of intensity of the XY data into regions of interest (ROI). Each ROI stores the intensity information over time as a trace. The traces were then distilled to extract the individual events. We used timescales of 1 to 256 s for the filtering during this process. Only oscillations with a z-score greater than 4 were considered as events. ROIs with an activity of below 5 oscillations over the whole experiments were discarded. Focusing on the dominant component, we filtered for events between 1 second and 10 seconds. For pharmacokinetics, we fitted a four-parameter logistic function. Kernel density estimations were calculated using a gaussian kernel with a bandwidth factor of 0.2.

### Network analysis

To create functional networks of the beta cells in islets, we assumed each ROI representing a node as part of an unweighted, undirected functional network graph. We calculated the cross-correlation between each ROI of an islet. Based on literature (reviewed in (20)) we used 0.7 as a threshold for the correlation to functionally connect the nodes. For each network obtained in this way, we calculated the mean node degree, the average clustering and the mean network efficiency during each condition to compare between the groups.

### Tomography of CR mouse beta cells and 3D reconstruction of ER compartments

Electron tomography was performed as previously described by us (15). Briefly, 300nm-thick sections of islets in situ were cut from pancreas blocks prepared for SEM with a Leica ultramicrotome and placed on 100-mesh copper grids. Tissue sections were coated with 20-nm colloidal gold particles and used as registration fiducials. Sample grids were imaged using a single-rotation tilt holder in a Tecnai High Base Titan (FEI; Hillsboro, OR) electron microscope operated at 300 kV. Grids were irradiated with electrons for ∼10 min to limit specimen thinning during imaging.

Sample illumination was held to near parallel and constant intensity beam conditions. Tilt series were captured using SerialEM software (University of Colorado, Boulder, CO) at a 0.81 nm per pixel resolution. A Gatan Ultrascan 4Kx4K CCD camera recorded the tomogram series. We collected 121 images per axis with 1 degree increment and a range of −60 to +60 degrees before repeating the acquisition protocol after rotating the sample holder by 90 degrees. We improved signal to noise ratio by applying a 2×2 pixel binning in IMOD (University of Colorado, Boulder, CO). The IMOD package with etomo java wrapper (https://en.wikipedia.org/wiki/IMOD) was used for alignment and registration of beta cell tomograms. Reconstruction of beta cell ER structures in the EM tomograms were achieved using WEBKNOSSOS (21), and the reconstructed volumes were exported as binary masks for downstream processing in Matlab for morphometric measurements.

### Analysis of mitochondria-ER contact landscapes

We ran individual beta cell SEM images through our data analysis pipeline as previously described (22). Briefly, cells were manually segmented using LabKit, and organelles were automatically segmented using trained UNet models to recognize ER or Mitochondria in SEM images, specifically. Next, a raster scan was performed to identify neighboring Mitochondria and ER pixels. Organelle contact sites were defined as areas at least 10nm (2px at 5nm resolution in the SEM images) in length in which organelle perimeters were within 10nm of each other. From these sites, mitochondria were defined as having ER contacts when >5% of their perimeter was part of an ER contact site.

### Statistics

Differences between the groups were calculated using either Mann-Whitney U test for non-normal, or Student’s t-test for normal distributed data in case of independent testing. Differences between distributions were assessed using the Kolmogorov-Smirnov test. For animal experiments and microscopy data, Student’s t test (Prism 9, GraphPad) was used to compare two groups, whereas a one-way ANOVA followed by Kruskal-Wallis test was used to compare two or more groups. A p value of <0.05 was considered statistically significant.

## Results

### Calorie restricted beta cells have altered cytosolic calcium activity

We have previously shown that beta cells from mice exposed to 2 months of CR sustain normoglycemia by secreting ∼50% less insulin *in vivo* than control AL mice (15). In that same study, isolated islet perifusion studies failed to find significant changes in insulin release between groups, which led us to hypothesize that removing islets from their native environment compromised normal beta cell function phenotypes. Therefore, we decided to investigate the cellular mechanisms underlying CR beta cell insulin secretion in acute live pancreatic slices. Importantly, slices maintain normal islet cell structure and function (19) without triggering stress responses that occur during the islet isolation process (23).

Acute live pancreatic slices from adult male FVB mice kept on AL or CR diet for 2 months were imaged for cytosolic [Ca^2+^] using high speed confocal microscopy. First, we established baseline and glucose-stimulated beta cell Ca^2+^ activity in AL and CR groups by imaging islets exposed to increasing glucose concentrations via a glucose ramp (from 6 mM to 8 mM to 10 mM glucose) (Figure 1A-B)). As expected, exposure to 6 mM conditions revealed that islets in both groups displayed limited oscillatory Ca^2+^ activity, which was significantly increased after exposure to 8 and even further to 10 mM glucose (Figure 1A-C). Notably, CR beta cells had a lower density of long-duration Ca^2+^ events defined by Ca^2+^ oscillatory event halfwidth and inter-event interval measurements (Fig. 1C-D). Moreover, this analysis revealed that CR beta cells show a significantly wider spectrum of Ca^2+^ event halfwidths, with a dominance of shorter events (Figure 1D-G). Specifically, fast Ca^2+^ oscillations in CR beta cells were ∼2.2 seconds faster at 6 mM glucose (AL: 4.04 s, CR: 2.84 s; p = 0.00031), ∼1.8 second faster at 8 mM glucose (AL: 5.36 s, CR: 3.41 s; p = 0.0001), and ∼1 seconds faster at 10 mM glucose treatment (AL: 4.24 s, CR: 3.21 s; p = 0.0001) (Fig. 1G, top pane).

**Figure 1.**
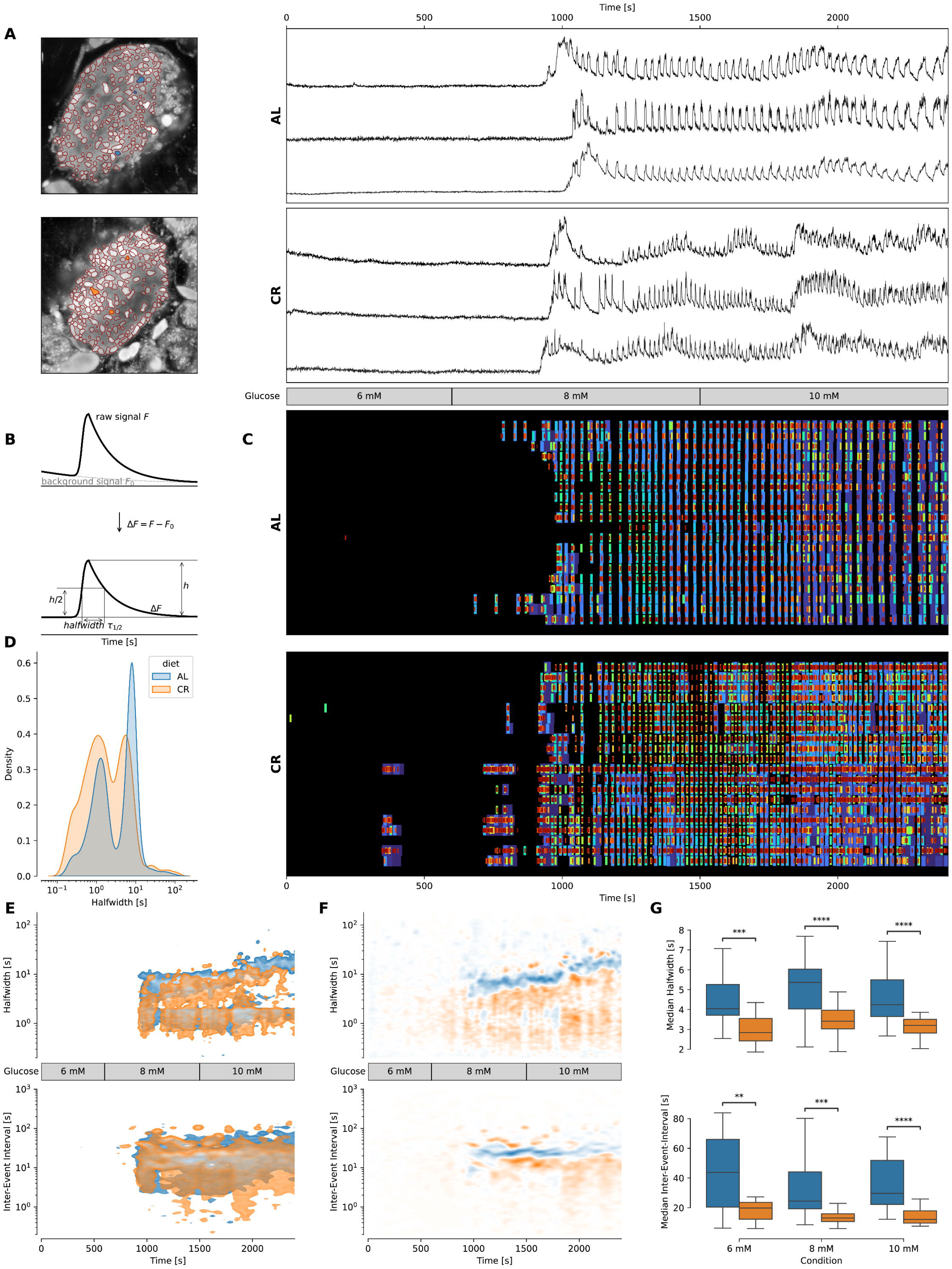
CR alters the Ca2+ response of beta cells during stimulating glucose conditions. **(A)** *Left*: Image of representative islets used for determining the traces. Colored ROIs indicate ROIs from which traces are plotted. *Right*: [Ca^2+^]_i_ traces (F/F_0_ to correct for bleaching artifacts) of a representative islet per group. The traces have been rebinned to show a frequency of 2 Hz. The treatment protocol is indicated in the middle and was the same for both groups. **(B)** Scheme illustrating quantification of event halfwidth. Raw signals were de-bleached. Then the full width of the half maximum amplitude was used as a robust readout. The time difference between the midtimes of sequent events were used to calculate the inter-event interval. **(C)** Raster plot showing events of the islets shown in (A) for 20 most representative ROIs per islet. Events are plotted over time, blue color indicates slower events, red color indicates faster events. **(D)** Kernel density estimation plot showing the density of halfwidths pooled over all experiments. It can be observed that the AL islets show more slow events, while CR islets tend to show more fast events.**(E)** *Top*: Kernel density estimation plots showing the halfwidth of the events pooled over all experiments with the stated protocol for each group. *Bottom*: Kernel density estimation plots Interevent interval for events of ROIs pooled over all experiments with the stated protocol. **(F)** *Top*: Kernel density estimation plots showing the difference of the KDEs plotted in (E, top pane) between both diet groups. *Bottom*: Kernel density estimation plots showing the difference of the KDEs plotted in (E, bottom pane). **(G)** Box plots showing the halfwidth (*top*) and median interevent-interval (*bottom*) of the events pooled over all experiments during the treatments indicated at the bottom for each group. Data obtained from AL islets are plotted in blue, data from CR islets in orange. Significance indications represent following p-values: * p < 0.05, ** p < 0.01, *** p < 0.001, **** p < 0.0001.

Next, we calculated the inter-event interval of Ca^2+^ events in AL and CR beta cells, which lead to the discovery of significant changes in the inter-event interval of the dominant fast events within the whole islet beta cell population. Specifically, inter-event intervals of fast Ca^2+^ oscillations in CR beta cells were ∼23 seconds shorter at 6 mM glucose (AL: 43.71 s, CR: 19.77 s; p = 0.0005), ∼11 second shorter at 8 mM glucose (AL: 24.40 s, CR: 13.00 s; p > 0.0001), and ∼17 seconds shorter at 10 mM glucose treatment (AL: 29.67 s, CR: 12.09 s; p < 0.0001)(Fig. 1G, bottom pane).

Together, these data indicate that CR beta cells have a higher frequency of faster Ca^2+^ events in response to stimulatory glucose concentrations, whereas AL beta cells have less and slower Ca^2+^ oscillatory events.

### CR beta cells have an elevated cAMP signaling

Previous imaging mass spectrometry experiments indicated that CR beta cells have higher concentration of cAMP(15). In beta cells, cAMP is a key second messenger involved in insulin exocytosis regulation by signaling via PKA (24) or Epac2 (18), to regulate ER Ca^2+^ release via IP_3_R and RYR2 (25), sensitivity of the secretory machinery to Ca^2+^ (26), while the effect on opening probability of voltage-activated Ca^2+^ channels (27) has been questioned (26). Therefore, we hypothesized that elevated levels [cAMP]_cyt_ levels in CR beta cells were responsible for modulating the observed CR beta cell phenotype (Figure 1).

To investigate differences in [cAMP]_cyt_ in AL and CR beta cells and how these could relate to Ca^2+^ homeostasis, we stimulated AL and CR slice islets with 6 or 8mM glucose together with a concentration ramp of epinephrine (Epi, 0.1 nM to 100 nM) (Figure 2A). This setup inhibits cAMP production and depletes cAMP signaling to suppress Ca^2+^-dependent insulin release via interruption of Ca^2+^ release from ER stores (28). As a result, CR beta cells would require higher concentrations of epinephrine to sustain the inhibition of Ca^2+^ oscillations due to maintenance of higher basal [cAMP]_cyt_ levels. As expected, CR beta cells had more frequent and shorter Ca^2+^ events under 8 mM glucose conditions (Figure 2A). Co-stimulation with 8 mM glucose and Epi caused significant changes to beta cell Ca^2+^ homeostasis including shortening of inter-event intervals (Figure 2B-C). Next, we calculated the minimum activity, maximum activity, EC_50_, and Hill coefficients for Epi stimulation in both groups by fitting the normalized inter-event interval against different concentrations of Epi in a four-parameter logistic function (Figure 2D). CR beta cells followed a distinctly different distribution compared to AL cells (! = 1.752 · 10^!”^, D = 0.37). CR beta cells had a 2-fold increase in Epi minimum frequency (AL: 0.0274 Hz, CR: 0.0448 Hz) and a slight increase in maximum frequency (AL: 0.0828 Hz, CR: 0.0931 Hz). Surprisingly, the EC_50_ for Epi was higher in the AL beta cells (AL: 11.415 nM, CR: 1.835 nM) (Figure 2B). These experiments support the idea that CR beta cells have higher cAMP signaling tone that supports faster glucose-stimulated Ca^2+^ oscillations.

**Figure 2.**
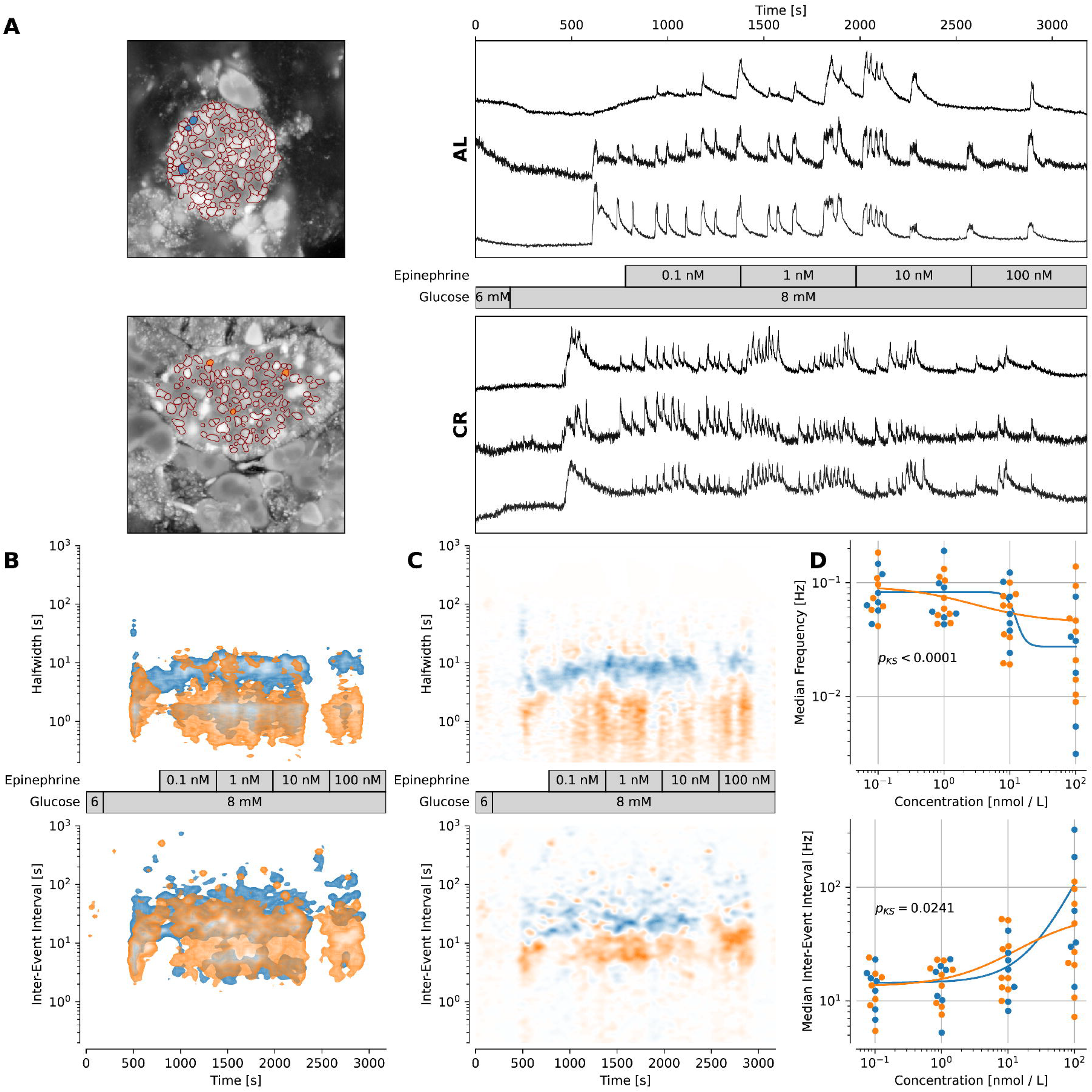
CR beta cells have higher [cAMP]_i_ signaling tone. **(A)** *Left:* Image of representative islets used for determining the traces. Blue/orange areas indicate ROIs, which were used to plot the traces. *Right:* [Ca^2+^]_i_ trace (F/F_0_ to correct for bleaching artifacts) of a representative islet per group. The traces have been rebinned to show a frequency of 2 Hz. The treatment protocol is indicated in the middle and was the same for both groups. **(B)** *Top:* Kernel density estimation plot showing the halfwidth of the events pooled over all experiments with the stated protocol for each group. *Bottom:* Kernel density estimation plot showing inter-event interval for events of ROIs pooled over all experiments with the stated protocol. **(C)** Difference of the respective kernel density estimation plots from (B). Blue areas indicate characteristics of the AL islets are dominant, while orange areas indicate dominant characteristics in CR islets. The protocol used is stated in the middle. **(D)** *Top:* Plot showing median frequency of events during epinephrine application for each used concentration. Dots indicate measured values. The Lines show a fitted four-parameter logistic function. P-value was calculated by using a Kolmogorov-Smirnov test to check for difference in distributions. *Bottom:* Plot showing median inter-event-interval during epinephrine application for each used concentration. Dots indicate measured values. Lines show a fitted four-parameter logistic function. P-value was calculated by using a Kolmogorov-Smirnov test to check for difference in distributions. Data obtained from AL islets are plotted in blue, data from CR islets in orange. Significance indications represent following p-values: * p < 0.05, ** p < 0.01, *** p < 0.001, **** p < 0.0001.

### Acetylcholine shifts the activity of AL beta cells into CR beta cell-like activity

ACh stimulates beta cell activity in a concentration-dependent manner by activating IP_3_Rs via M3-receptor coupled to Gq protein, PLC activation and production of IP_3_R (29). We hypothesized that an increased tonus of IP_3_ in AL beta cells should mimic the phenotype of the CR beta cells with fast oscillation frequency and reduced inter-event intervals. In fact, application of the saturating ACh concentration significantly reduced inter-event interval(AL: [8 mM glucose: 17.89 s, ACh: 9.08 s], CR: [8 mM glucose: 11.22 s, ACh: 7.99 s]) and median halfwidth of the events (AL: [8 mM glucose: 4.42 s, ACh: 2.43 s], CR: [8 mM glucose: 3.79 s, ACh: 2.62 s]) (Figure 3D). After removal of ACh the inter-event interval has been partially restored towards longer values in AL beta cells (AL: 9.23 s, CR: 7.98 s). Taken together, in CR beta cells increased [cAMP]_cyt_ enables cells to operate with higher IP_3_R activity.

**Figure 3.**
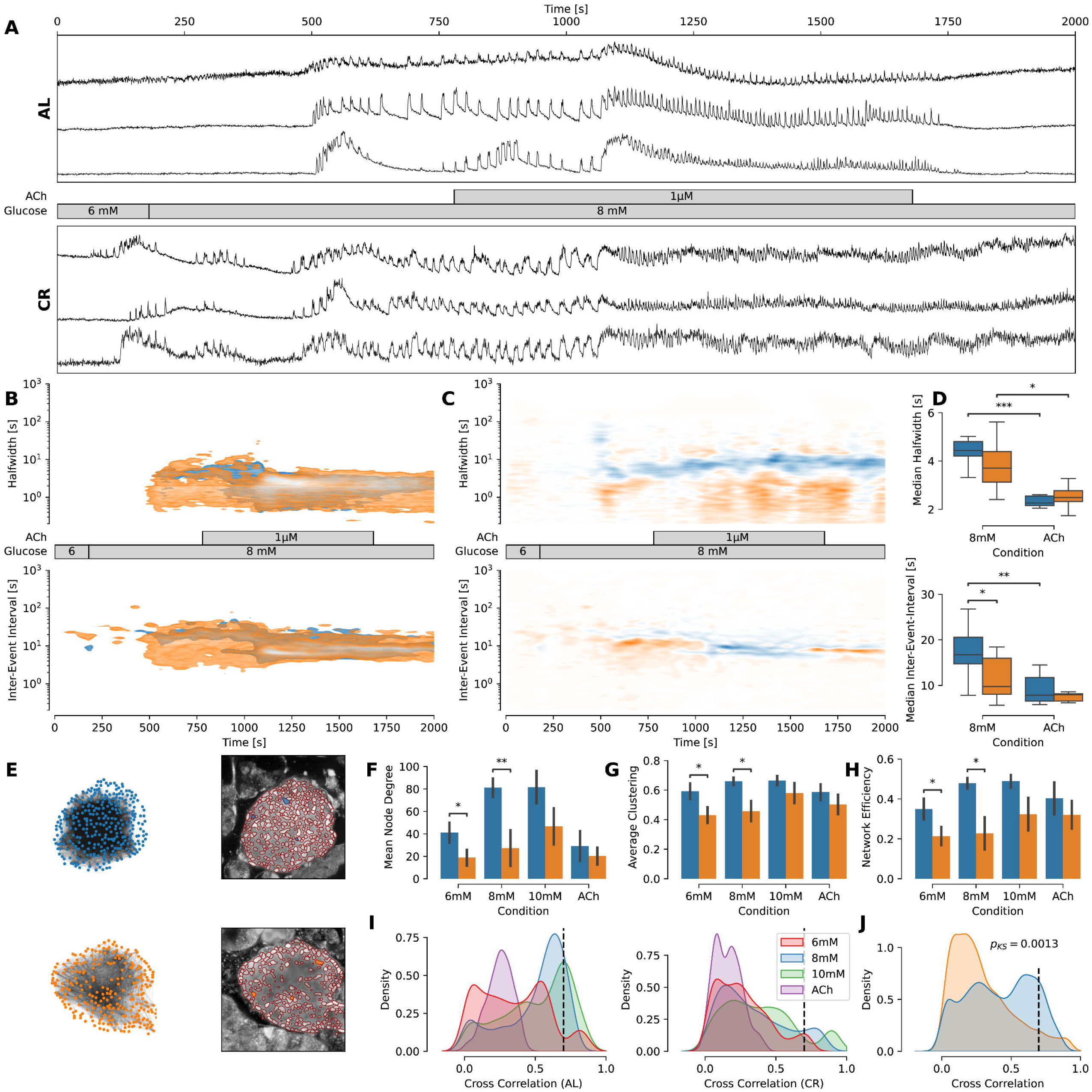
CR beta cells have increased resistance to acute ER stress and lose network connectivity. **(A)** [Ca^2+^]_i_ trace (F/F_0_ to correct for bleaching artifacts) of a representative islet per group. The traces have been rebinned to show a frequency of 2 Hz. The treatment protocol is indicated in the middle and was the same for both groups. **(B)** *Top*: Kernel density estimation plot showing the halfwidth of the events pooled over all experiments with the stated protocol for each group. *Bottom*: Kernel density estimation plot showing inter-event interval for events of ROIs pooled over all experiments with the stated protocol. **(C)** Difference of the respective kernel density estimation plots from (C). Blue areas indicate characteristics of the AL islets are dominant, while orange areas indicate dominant characteristics in CR islets. The protocol used is stated in the middle. **(D)** Box plot showing the change of median inter-event interval and median halfwidth during the treatment change from 8 mM glucose to 1 µM ACh. **(E)** *Left*: Functional network plotted from the islet shown in panel (A). Dots indicate nodes, lines indicate edges. Nodes were connected if the correlation between the traces exceeded at least a value of 0.7 to explain ∼50 % (R^2^ > 0.49) of the data. *Right*: Microscopical images of respective islets with ROIs used to measure traces. Highlighted ROIs are representative for the islets’ activity. **(F-H)** Network parameters observed per islet pooled for each condition. Bars indicate mean values ± SEM. **(I)** KDE plots showing the distribution of cross correlations for each group respectively colored by the condition. The dashed line indicates the 0.7 cut-off threshold being used to connect two nodes in the functional network. **(J)** KDE plots showing the distribution of cross correlations between each group across all conditions. The dashed line indicates the 0.7 cut-off threshold being used to connect two nodes in the functional network. P-value was determined by KolmogorovSmirnov test between both distributions. Data obtained from AL islets are plotted in blue, data from CR islets in orange. Significance indications represent following p-values: * p < 0.05, ** p < 0.01, *** p < 0.001.

### Acetylcholine-induced stress in CR beta cell collectives leads to higher resistance to depletion of ER Ca^2+^ stores

[cAMP]_cyt_ drives phosphorylation of RyRs and IP_3_Rs to regulate ER Ca^2+^ flux (30). Here, we investigated the hypothesis that CR-induced changes in beta cell Ca^2+^ oscillations were due to modulation of ER Ca^2+^ homeostasis mechanisms. To test this hypothesis, we used a supraphysiological ACh (1 µM) stimulation of tissue slices after stimulation with 8 mM glucose to activate beta cell IP_3_Rs and trigger ER Ca^2+^ efflux.

As expected, ACh stimulation led to oscillatory Ca^2+^ responses in beta cells from both diet groups that decayed overtime (Figure 3C-E). Interestingly, we find that both event median halfwidth (before ACh: 4.42 s, after ACh: 2.43 s, ! = 0.0003) and median inter-event interval (before ACh: 17.89 s, after ACh: 9.08 s, ! = 0.0028) are significantly decreased in AL beta cells, whereas in CR beta cells only median halfwidth is decreased after ACh treatment (before ACh: 3.78 s, after ACh: 2.62 s, ! = 0.0217). This indicates CR islets being able to run individual stimuli longer even in stressed conditions.

### CR decreases beta cell network connectivity

Several studies by us and others have established the spatial and functional relationship between neighboring beta cells within individual islets (31; 32). Importantly, beta cells have coordinated Ca^2+^ activity patterns mediated by gap junctions (33) that create interconnected beta cell networks (20). Glucose stimulation increases beta cell network connectivity to potentiate beta cell insulin release and control glucose homeostasis (34; 35). Therefore, we hypothesized that the reduced insulin secretion of CR beta cells (15) correlated to loss of beta cell connectivity between in CR islets.

To test this hypothesis, we calculated the functional relationship of each individual beta cells in AL and CR islets under basal and rising glucose concentrations and ACh stimulation (Figure 3E). We observed expected changes in cross-correlations for each treatment within the groups (Figure 3I), and significant differences overall showing more abundant cross-correlation values in AL beta cells (! = 0.0013, Figure 3J). To reconstruct and quantify beta cell networks, we used three separate indexes such as *mean node degree* (number of connections between beta cells), *average clustering* (the probability that two beta cell neighbors are also connected to each other), and *network efficiency* (how many connections are needed to connect one end of the network to another). As expected, during basal glucose conditions and glucose stimulation we could observe a significant increase in beta cell connectivity (6 mM: [AL: 41.05, CR: 18.84, ! = 0.04], 8 mM: [AL: 81.24, CR: 27.35, ! = 0.0097]), clustering (6 mM: [AL: 0.59, CR: 0.43, ! = 0.03], 8 mM: [AL: 0.66, CR: 0.46, ! = 0.0152]), and network efficiency (6 mM: [AL: 0.35, CR: 0.21, ! = 0.0397], 8 mM: [AL: 0.48, CR: 0.23, ! = 0.0118]) in AL mice (Figure 3F-H), indicating that CR beta cell networks are significantly suppressed (or disconnected) under physiological conditions.

Together, these results indicate that the CR islet beta cell population loose strong connectivity favoring weaker interactions (36). This could allow finer control of insulin release and faster adaptation to gradual decreases in insulin demand due to increased insulin sensitivity in CR mice (15).

### CR beta cells have normal ER architecture

In post-mitotic cells, CR significantly alters ER Ca^2+^ homeostasis by elevating expression of Ca^2+^-buffering chaperones (e.g., CalR) (37). The data presented here (Figures 1-3) indicates that CR beta cells have increased ER Ca^2+^ activity, which could be potentially explained by changes in ER ultrastructure (i.e., larger ER Ca^2+^ storing capacity) or by changes in ER-mitochondria contacts that can mediate Ca^2+^ transfer between organelles (38). To investigate how CR impacts the structure of the beta cell ER *in situ*, we analyzed our previously published beta cell electron microscopy datasets acquired using scanning electron microscopy (SEM) or using electron tomography (eTomo) (15) and analyzed these images using applied machine learning-based organelle segmentation and connectome analysis (recently established by us (22)). This allowed us to quantify the density of mitochondria and ER contacts in AL and CR beta cells and revealed that most beta cell mitochondria (∼95%) are uncoupled from ER structures (Figure 4A-B). In fact, most ER and mitochondria are located within 20-to-50nm away from their nearest point (Figure 4A-B). However, no differences in mitochondria-ER contacts were found between AL and CR beta cells were found, thus indicating that contact by these two organelles is not modulated by CR. Likewise, and using eTomo, we found no changes to beta cell ER size and volume in the reconstructed 3D ER volumes (Figure 4C-E).

**Figure 4.**
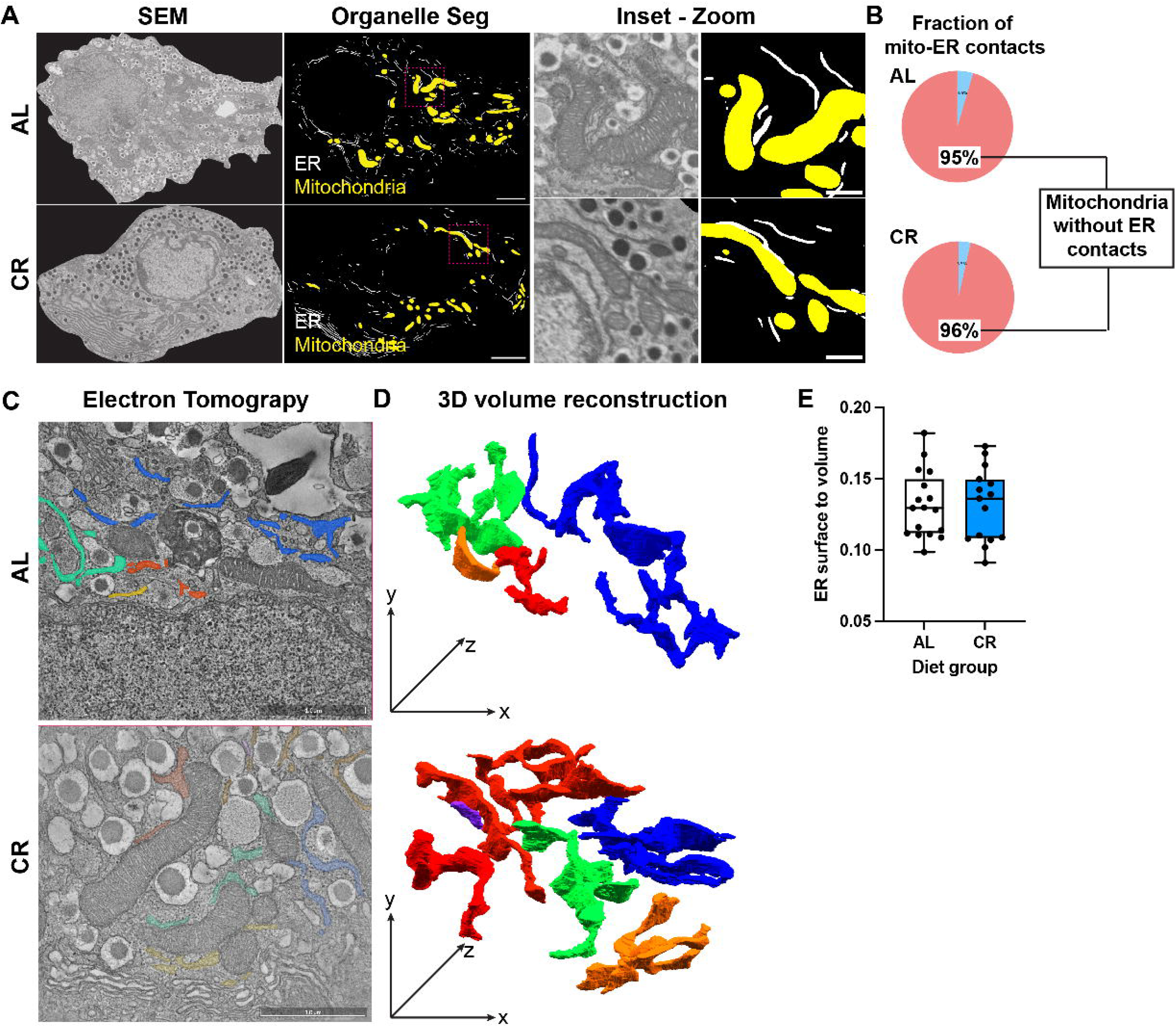
CR does not alter beta cell ER structure or mitochondria-ER contact density. **(A)** Representative scanning electron microscopy (SEM) and mitochondria (yellow) and ER (white) organelles segmented using trained 2D U-nets applied to SEM data. Inset, SEM micrographs revealing the location of Mitochondria-ER contact sites. Magenta arrows point to mitochondria-ER contact sites. **(B)** Relative fraction of detected mitochondria-ER contacts in AL and CR beta cells. **(C)** Representative electron tomography micrographs of beta cells from AL and CR mice, with mitochondria, lysosomes, insulin granules, golgi, and nucleus in view. Yellow, blue, green, and orange objects mark the location of reconstructed ER structures. **(D)** Representative 3D reconstruction of ER structures in the tomograms showing a network-like structure of ER complexes. **(E)** Quantification of the surface area to volume ratio of reconstructed ER structures from AL and CR beta cells. Each dot represents an individual ER object.

Therefore, this data demonstrates that short term CR of young adult mice does not impact the structure of beta cell ER nor the density of mitochondria-ER contact sites and thus indicate that the observed Ca2+ homeostasis phenotype of CR beta cells is likely due to changes in mechanisms regulating Ca2+ movement, such as ion channels IP3R and RyRs (Figures 2-3).

## Discussion

In this study we investigated the Ca^2+^ oscillation patterns of beta cells from AL and CR mice under basal, glucose-stimulated, [cAMP]_cyt_ depleted and high IP_3_ load conditions. Using high speed confocal microscopy of acute pancreatic tissue slices and mathematical modeling, we demonstrate that CR beta cells operate with shorter events with shorter inter-event intervals that correlate with an overall collapse of beta cell functional network architecture. The AL beta cells can be forced into a similar phenotype with high IP_3_ load following ACh stimulation, however they still exhaust sooner than CR beta cells. These findings are not explained by changes to overall ER architecture, thus suggesting that likely changes in Ca^2+^ flux at the ER level and its interactions with [cAMP]_cyt_, explain the changes in Ca^2+^ homeostasis of CR beta cells. Moreover, we propose the functional reorganization of beta cell Ca^2+^ flux underlies the markedly reduced levels of beta cell insulin secretion observed during CR, which is sufficient to maintain normoglycemia due to a higher level of peripheral insulin sensitivity (15).

The exact relationship between Ca^2+^ oscillations and insulin secretion has been elusive for high frequency recordings (39), possibly since these aspects have been followed using dissociated beta cells or isolated islets that typically do not recapitulate the complexity of in situ Ca^2+^ homeostasis. Our results demonstrate that islet beta cells from acute tissue slices of CR mice have a reduced overall Ca^2+^ response when stimulated by glucose or ACh versus AL islets, which in turn have longer Ca^2+^ transients (Figure 1). The question is, what drives the changes in the duration of events and the intervals between them? The Ca^2+^ flux contributing to [Ca^2+^]_cyt_ is a consequence of a delicate interplay between major transport proteins in pancreatic beta cells, like ATP-driven SERCA pumps (for extrusion of Ca^2+^ from the cytosol into the ER), and VDCCs, store-operated Ca^2+^ entry (SOCE) channels, RyR1-3, IP_3_Rs and other cationic channels (which transport Ca^2+^ into the cytosol). The first explanation for the described changes in the [Ca^2+^]_cyt_ kinetics would center on VDCCs, which are well described as key regulators of Ca^2+^ events in cardiomyocytes, where cAMP-dependent activation of both VDCCs and SERCA kinetics support both inotropy and reduced interval between contractions. There are two arguments speaking against the role of VDCCs in beta cells. Our own patch-clamp experiments could not reproduce cAMP-dependent increase in opening probability (26) and inability of selective blockers of L-type Ca^2+^ channels in the ECS to prevent glucose-dependent activation of beta cells in tissue slices (5).

To explain shorter event halfwidth and the shorter inter-event interval in CR beta cells we propose the dominance of the ER Ca^2+^ flux after CR diet. We have no molecular evidence for the presence of a SERCA modulator, like phospholamban in beta cells. Due to availability of physiological ligands, we focused in this study on the role of IP_3_R stimulation, however, at this stage we cannot estimate or exclude the contribution of RyRs of being able to contribute to the faster [Ca^2+^]_cyt_ kinetics in CR beta cells. As we see in Fig 3., ACh reversibly stimulates both AL beta cells to significantly shorter events with shorter inter-event intervals on a cell collective level in comparison to 8 mM glucose stimulation only. In CR beta cells, ACh did not significantly change the inter-event interval.

We have previously shown that CR beta cells have more mitochondria with increased mitochondrial cristae density, and CR islets have elevated [cAMP]_cyt_ (15). Together, these suggest CR beta cells have higher metabolism resulting in higher [ATP] and [cAMP]_cyt_, which would lead to elevated tone of ER Ca^2+^ homeostasis due to high activity of SERCA, IP_3_Rs and RyRs. The resulting increased leak current could be the reason why we see more events in the fast time domain in the CR beta cells. In the present study, we show that the reaction of CR islets to glucose differs compared to AL islets: CR islets react with more events, and shorter events in the fast time domain, which reflects the reported higher [cAMP]_cyt_. To further substantiate our hypothesis, we decided to first check the effect of pharmacologically decreased [cAMP]_cyt_ levels. In fact, CR beta cells exposed to epinephrine (which depletes cAMP) can sustain Ca^2+^ transients for longer periods of time, thus supporting the idea that CR beta cells mice have higher basal [cAMP]_cyt_ levels to support the enhanced activity of the IP_3_R_s_. Moreover, we mimicked elevated IP_3_ conditions using high levels of ACh to strongly stimulate IP_3_R-mediated Ca^2+^ release from the ER to test how long the endogenous ER Ca^2+^ and found that CR beta cells can sustain a high Ca^2+^ homeostasis for longer periods of time. Together, these experiments support our previous findings and show that CR increases beta cell [cAMP]_cyt_ to regulate ER Ca^2+^ fluxes. The exact mechanisms that increase cAMP synthesis in CR beta cells remain unknown.

Finally, CR beta cells are largely post-mitotic and are thought to be more mature due to increased expression of beta cell identity genes (15). Because of this phenotype, we assumed that CR islets would be more functionally connected, however CR beta cells formed largely disconnected networks where the activity of each individual cell was weakly correlated to other beta cells in an islet. These findings hint that longer-lived beta cells may not rely on strong functional connectivity to maintain glucose homeostasis to coordinate insulin release.

Putting everything together, we show that calorie restriction might not only influence the beta cells of mice on a morphological level, but also on a functional level. This study provides further evidence that [cAMP]_cyt_ in islets of CR mice is higher than in AL mice. Furthermore, CR mice can control and maintain beta cell functionality with higher speed and precision. We show that this is due to better Ca^2+^ homeostasis in the ER either by higher [Ca^2+^]_ER_ or changes in channel properties.

However, the exact mechanism is not known, providing opportunities for future experiments to explore this more deeply. Importantly, this study characterizes the impact of calorie restriction on long-lived beta cell Ca2+ homeostasis and provides a mechanistic explanation on how postmitotic beta cells adapt when the metabolic demand for insulin release is reduced due to increased peripheral insulin sensitivity.

## Author contributions

J.P. conducted all Ca^2+^ imaging experiments, performed data analyses, created figures and wrote the manuscript. CDS, M.C., C.A., A.H., A.C., G.P., and M.H.E. collected data and performed image analysis. R.A.D. and M.S.R. conceptualized the project, provided the analytical pipeline for imaging analysis, performed data analyses, and wrote the manuscript.

## Acknowledgments

his research was supported by recruitment funds from the Vanderbilt’s Department of Molecular Physiology and Biophysics and NIH-NIDDK grant R01DK138141 to RAeD, grants by the Austrian Science Fund/Fonds zur Förderung der Wissenschaftlichen Forschung (bilateral grants I3562-B27 and I4319-B30), a grant from Vienna Science and Technology Fund WWTF (LS23-026), from NIH (R01DK127236), and from the Slovenian Research Agency (research core funding program P3-0396). M.H.E. is supported by the NIH-NINDS grant U24NS120055.

## Notes

### Competing Interest Statement

The authors have declared no competing interest.

